# The Role of Microglia and Complement C5/C5a in the Pathogenesis of Rhegmatogenous Retinal Detachment with Choroidal Detachment

**DOI:** 10.1101/2025.02.15.637446

**Authors:** Huiyan Xu, Qiuhong Wang, Xuan Chen, Qingyu Huang, Shasha Xu, Zhifeng Wu

## Abstract

**Background:** Rhegmatogenous retinal detachment with choroidal detachment (RRDCD) is an uncommon and sight-threatening disorder marked by fast development and significant inflammation. This study aimed to identify cellular and molecular signatures distinguishing RRDCD from typical rhegmatogenous retinal detachment (RRD) and to investigate the roles of microglia and the complement C5/C5a pathway in disease pathogenesis.

**Methods:** Single-cell RNA sequencing (scRNA-seq) was employed to analyze vitreous samples from patients with RRD and RRDCD, indicating its involvement in blood-retina barrier impairment.

**Results:** Our findings revealed a distinct cellular landscape in RRDCD, characterized by enhanced connectivity between microglia and dendritic cells, alongside a significant upregulation of the complement C5-C5AR1 interaction. In vitro experiments indicated that treatment with complement C5 promoted microglial proliferation and activation, induced apoptosis in RF/6A endothelial cells, and disrupted tight junctions in ARPE-19 epithelial cells, suggesting a role in blood-retina barrier dysfunction.

**Conclusion:** The findings substantiate the inflammatory hypothesis regarding the pathogenesis of RRDCD, emphasizing the critical functions of microglia and the complement C5/C5a pathway in intensifying retinal inflammation and undermining vascular integrity.

## Introduction

Rhegmatogenous retinal detachment with choroid detachment (RRDCD) is a complex and uncommon type of rhegmatogenous retinal detachment (RRD), distinguished by fast advancement and significant inflammatory reactions. This disorder frequently requires vitrectomy and is linked to a high postoperative recurrence rate, resulting in considerable vision impairment. The prevalence of RRDCD differs worldwide, with significantly elevated rates observed in China^[1–3]^. Notwithstanding its clinical importance, the pathophysiology of RRDCD is inadequately comprehended, hindering the formulation of effective therapeutic methods.

RRDCD is a vision-threatening disorder that differs from standard RRD due to the involvement of the choroid, a vascular layer located between the retina and the sclera. The presence of choroidal detachment in RRDCD signifies a broader disruption of ocular tissues and a possibly more aggressive disease progression. This condition is often associated with a pronounced inflammatory response, which may result in proliferative vitreoretinopathy (PVR)—a complication that considerably impedes the efficacy of surgical treatment.^[4,5]^.

Our previous studies^[6–9]^ have indicated that various cytokines and inflammatory-related proteins, including components of the complement system such as C1r, C3, C5, C6, and C8, are significantly expressed in the vitreous of patients with RRDCD compared to those with RRD. The complement system, a part of the innate immune system, plays a critical role in immune surveillance and inflammation. Its dysregulation has been implicated in a variety of ocular diseases, including age-related macular degeneration and retinitis pigmentosa.

The specific cellular players involved in the pathogenesis of RRDCD and their interactions with the complement system have not been fully elucidated. Microglia, the resident immune cells of the central nervous system—including the retina—have been suggested to contribute to retinal inflammation and homeostasis. They are known to respond to environmental changes, such as those occurring during retinal detachment, and can influence disease progression through mechanisms including phagocytosis and cytokine production^[10–12]^.

Given the complexity of RRDCD and the potential role of microglia and the complement system in its pathogenesis, a deeper understanding of the cellular and molecular mechanisms involved is crucial. Single-cell RNA sequencing (scRNA-seq) ^[13]^offers a powerful tool to dissect the heterogeneity of cell populations in vitreous samples and identify specific cell types and pathways that may serve as therapeutic targets.

In this study, we employed scRNA-seq to investigate the cellular landscape in vitreous samples from patients with RRD and RRDCD, with a particular focus on microglia and the complement C5/C5a pathway. Our aim was to uncover the cellular and molecular signatures that distinguish RRDCD from RRD and to explore the potential roles of microglia and complement C5/C5a in disease pathogenesis. Such insights could lead to new therapeutic strategies for this devastating condition.

## Materials and Methods

### Patient Selection

Patients diagnosed with primary RRD or RRDCD, without other ocular diseases as determined by clinical evaluations and laboratory examinations, were enrolled between January 2021 and March 2023. Exclusion criteria included vitreous hemorrhage, ocular trauma, glaucoma, previous intraocular surgery, ocular tumors, and significant ocular or systemic diseases.

### Single-Cell RNA Sequencing (scRNA-seq)

#### Sample Collection and Preparation

Vitreous samples were obtained during 25-gauge pars plana vitrectomy performed by a single experienced surgeon. Approximately 3 mL of vitreous fluid was collected near the vitrectomy entry site from each patient and immediately placed on ice for subsequent processing.

#### Library Preparation and Sequencing

Using the 10× Genomics Chromium Single Cell 3’ protocol, individual cells were partitioned into nanoliter-scale droplets containing gel beads. Within each droplet, cell lysis and reverse transcription occurred, yielding cDNA molecules sharing a unique 10× barcode per cell. This barcoding facilitated the separate indexing of each cell’s transcriptome. Following barcoding, cDNA libraries were constructed and sequenced using Illumina platforms, generating paired-end reads corresponding to the 5′ and 3′ ends of the cDNA fragments.

#### Data Processing and Quality Control

Sequencing data were processed using Cell Ranger (10× Genomics) for initial alignment, barcode processing, and gene expression quantification. Subsequent data analysis was conducted using Seurat, an R package designed for the analysis and visualization of single-cell RNA-seq data. Stringent quality control measures were implemented to ensure data integrity and reliability, including filtering out low-quality cells, cells with high mitochondrial gene expression, and genes expressed in fewer than a minimum number of cells.

#### Dimensionality Reduction and Clustering

High-dimensional scRNA-seq data were subjected to Principal Component Analysis (PCA) to identify significant sources of variation. Uniform Manifold Approximation and Projection (UMAP) was used to map the data into two-dimensional space for clearer interpretation. Clustering algorithms such as Louvain were applied to the PCA-reduced data to group cells with similar gene expression profiles.

#### Cell-Cell Communication Analysis

To explore cell–cell communication patterns, we employed CellPhoneDB, a database designed for analyzing ligand–receptor interactions. This tool enabled the identification of ligand–receptor pairs and their expression across different cell types, providing insights into the active signaling pathways within the cellular microenvironment.

### Primary Mouse Retinal Microglia Isolation and Identification

#### Isolation Procedure

Primary retinal microglia were isolated from 1-3 day-old neonatal mice. Culture plates were precoated with poly-D-lysine (50 µg/ml) in DMEM/F12 the day prior to extraction. Mice were euthanized via intraperitoneal injection of 0.5 ml of 10% chloral hydrate. Under aseptic conditions, eyes were carefully excised using sterile scissors and forceps and immediately placed in phosphate-buffered saline (PBS). A central window in the cornea was created using a syringe, and ophthalmic scissors were used to incise along the limbus. The choroid was gently detached, and structures such as the lens, vitreous body, and sclera were removed, retaining only the retina and choroid. The isolated tissues were transferred to a centrifuge tube containing trypsin digestion solution and mechanically dissociated into fragments. The mixture was incubated at 37°C for 25 minutes, after which DMEM/F12 supplemented with 10% fetal bovine serum (FBS) was added to neutralize the trypsin. The cell suspension was passed through a 75 µm filter to remove undigated tissue and centrifuged. The poly-D-coated plates were washed three times with PBS to eliminate residual matrix. The cell pellet was resuspended in DMEM/F12 containing 10% FBS and 1% penicillin-streptomycin and seeded into culture flasks. Cells were incubated overnight at 37°C with 5% COL to allow attachment. The following day, cell morphology, density, adherence, and growth were assessed under a microscope, and the medium was refreshed as necessary.

#### Identification of Microglia

Isolated microglia were identified using immunocytochemical staining for ionized calcium-binding adapter molecule 1 (IBA-1), a specific marker for microglia in the central nervous system. Cells were fixed with 4% paraformaldehyde, permeabilized, and incubated with an anti-IBA-1 primary antibody, followed by a fluorescently labeled secondary antibody. Nuclei were counterstained with DAPI.

### Complement C5 Treatment and Co-Culture Systems

#### Treatment Protocol

Microglia were divided into experimental and control groups. Experimental groups were treated with complement C5 at concentrations of 0.5 µg/ml, 1.0 µg/ml, and 2.5 µg/ml, while the control group received no complement C5.

#### Co-Culture with RF/6A Endothelial Cells

For co-culture with RF/6A endothelial cells, microglia were seeded in the upper chamber of Transwell inserts and allowed to adhere for 2 hours. Non-adherent cells were gently removed with PBS. Different concentrations of C5 were added to the DMEM medium in the upper chamber, and RF/6A cells were seeded in the lower chamber. The co-culture system was incubated at 37°C with 5% COL for 24 hours. Cell viability of RF/6A cells was assessed using the MTT assay, and apoptosis was evaluated using the TUNEL assay. Based on these results, a C5 concentration of 1.0 µg/ml was selected for further experiments.

#### Co-Culture with ARPE-19 Epithelial Cells

For co-culture with ARPE-19 epithelial cells, microglia treated with 1.0 µg/ml C5 were seeded in the upper chamber of Transwell inserts and allowed to adhere for 2 hours. Non-adherent cells were removed with PBS, and ARPE-19 cells were seeded in the lower chamber. The co-culture was maintained for 24 hours, after which the expression of the tight junction protein ZO-1 in ARPE-19 cells was examined using immunofluorescence staining.

### Cellular Assays

#### Cell Viability Assay

Cell viability was assessed using the MTT assay. Microglial cell suspensions were adjusted to a density of 10,000 cells per well and inoculated into 96-well plates. Edge wells were filled with sterile PBS to prevent evaporation effects. After overnight incubation at 37°C with 5% COL to form a confluent monolayer, cells were treated with different concentrations of C5 (0 µg/ml or 1.0 µg/ml) by adding 100 µl of each C5 concentration to the respective wells, with six replicates per treatment. After 24 hours of treatment, cell morphology was observed under an inverted microscope. Subsequently, 20 µl of 0.5% MTT solution was added to each well and incubated for 2 hours at 37°C. The culture medium was removed, and 150 µl of dimethyl sulfoxide (DMSO) was added to dissolve the formazan crystals. Absorbance was measured at 490 nm using a microplate reader to determine cell viability.

#### Immunofluorescence Detection

Microglia were seeded in 24-well plates and treated with 1.0 µg/ml C5 or left untreated as controls. After 24 hours of incubation, cells were washed twice with PBS and fixed with 4% paraformaldehyde for 15 minutes. Cells were permeabilized and blocked with blocking buffer for 1 hour. Primary antibodies were applied and incubated overnight at 4°C. The following day, cells were washed and incubated with fluorescently labeled secondary antibodies for 1 hour at room temperature. After three washes, nuclei were stained with DAPI. Fluorescence microscopy was used to observe and document the staining patterns.

#### Western Blot Analysis

Protein extraction was performed by washing cells three times with cold PBS followed by digestion with trypsin. Cells were collected into centrifuge tubes, centrifuged to remove the supernatant, and washed twice with cold PBS. Protein was extracted by adding RIPA buffer containing PMSF and thoroughly homogenized. Lysates were incubated on ice for 30 minutes and then centrifuged at 12,000 rpm for 15 minutes at 4°C. The supernatant containing proteins was collected and stored at -80°C. Proteins were denatured by adding loading buffer and boiling at 100°C for 5-10 minutes. Equal amounts of protein samples were loaded into SDS-PAGE gels and separated at 80V for 20 minutes (stacking gel) followed by 100V for 60 minutes (separating gel). Proteins were then transferred to PVDF membranes at 400 mA for 40 minutes. Membranes were blocked with 5% non-fat milk in TBST to prevent non-specific binding, followed by incubation with primary antibodies overnight at 4°C. After washing, membranes were incubated with fluorescently labeled secondary antibodies. Protein bands were visualized using a chemiluminescence imaging system.

#### Enzyme-Linked Immunosorbent Assay (ELISA)

After 24 hours of C5 treatment, supernatants from microglia cultures were collected and subjected to ELISA to quantify the levels of pro-inflammatory cytokines IL-1β and TNF-α. The assays were performed according to the manufacturer’s protocols. Briefly, 100 µl of each sample was added to ELISA plates pre-coated with capture antibodies specific for IL-1β and TNF-α. Following incubation and washing, detection antibodies and corresponding substrates were added. Absorbance was measured at 450 nm, and cytokine concentrations were determined using standard curves.

#### TUNEL Assay for Apoptosis Detection

Apoptosis in RF/6A endothelial cells was assessed using the TUNEL assay. Cells were fixed with 4% neutral paraformaldehyde for 10 minutes at room temperature. Fixed cells were then treated with the TUNEL reaction mixture according to the manufacturer’s instructions. Subsequently, cell suspensions were applied to glass slides and allowed to dry. Slides were washed twice with PBS for 5 minutes each. TUNEL-positive nuclei indicating apoptotic cells were visualized and quantified under a fluorescence microscope.

## Statistical Analysis

All experiments were performed in triplicate and repeated independently at least three times. Data are presented as mean ± standard deviation (SD). Statistical significance was determined using one-way analysis of variance (ANOVA) followed by Tukey’s post hoc test. A p-value of less than 0.05 was considered statistically significant. Analyses were conducted using R software (version 4.4.2) or GraphPad Prism (version 10.0).

## Result

### Patient Demographics and Clinical Characteristics

Single-cell transcriptome analysis was conducted on samples collected from four patients diagnosed with RRD and four patients diagnosed with RRDCD. Detailed information about patient demographics and clinical characteristics is provided in Table 1.

**Table 1.** Baseline data for the three patients with RRD and RRDCD in the study.

| Group | Sex | Age | Course<br>(days) | Visual acuity | IOP (mmHg) | Severity |
| --- | --- | --- | --- | --- | --- | --- |
| RRD1 | Female | 52 | 6 | OD0.4 OS0.2 | OD13.5 OS14.4 | OS PVRB, RD in two quadrants |
| RRD2 | Male | 64 | 21 | OD0.06 OS0.5 | OD10.3 OS10.9 | OD PVRB, RD in two quadrants |
| RRD3 | Male | 51 | 5 | OD0.4 OS0.25 | OD12.3 OS8.3 | OS PVRB, RD in two quadrants |
| RRD4 | Male | 62 | 5 | OD0.8 OS0.4 | OD9 OS6.6 | OS PVRB, RD in one quadrant |
| RRCD1 | Male | 44 | 10 | OD0.3 OS0.4 | OD7 OS13.2 | OD PVRB, RD+CD in one quadrant |
| RRCD2 | Male | 48 | 7 | OD CF OS0.5 | OD6.8 OS13.3 | OD PVRB, RD in two quadrants, ciliary choroid detachment in 360 degrees |
| RRCD3 | Female | 49 | 5 | OD HM OS NLP | OD5.2 OS26.1 | OD PVRB, RD+CD in two quadrants |
| RRCD4 | Female | 63 | 7 | OD hand move OS1.0 | OD6.8 OS16.4 | OD PVRB, RD+CD in two quadrants |

Figure 1 illustrates the findings from B-scan ultrasound and ultrasound biomicroscopy (UBM) of the patients. In RRD patients, B-scan ultrasound revealed retinal detachment, while UBM showed no detachment of the ciliary body or choroid. In contrast, RRDCD patients exhibited retinal detachment or accompanying choroidal detachment on B-scan ultrasound, and UBM revealed associated detachment of the ciliary body and choroid.

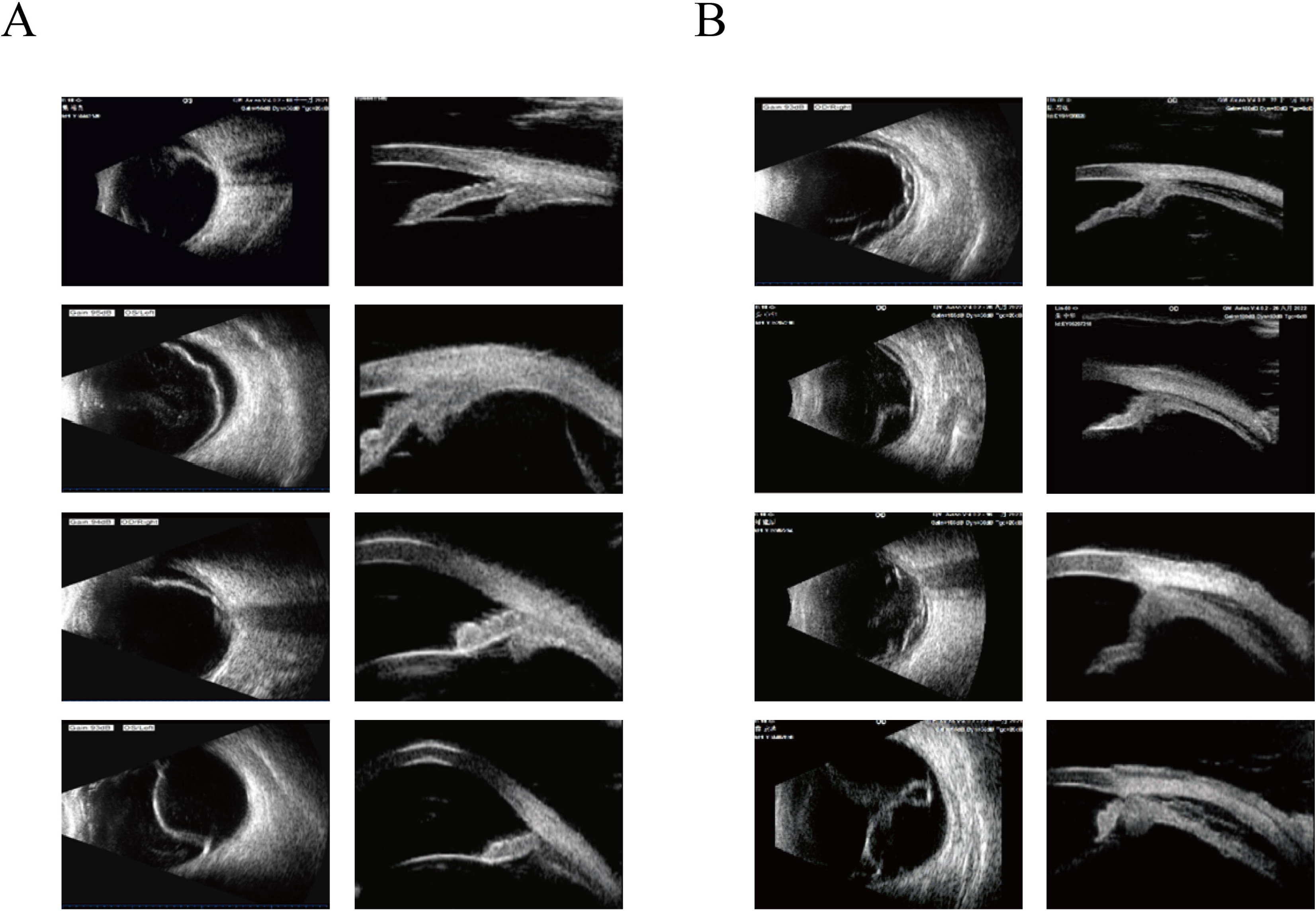

### Clustering and Cellular Composition

We performed scRNA-seq to investigate cellular heterogeneity in RRD and RRDCD patients. The results of dimensionality reduction, clustering, and cell type annotation are presented in Figure 2. Utilizing UMAP effectively visualized distinct cellular populations from both patient groups. Cells were initially segregated into immune and non-immune populations based on the expression of the immune cell marker CD45 (PTPRC). Subsequent cell type annotation was conducted using the CellMarker 2.0 database and previous studies. Figure 2B’s bubble plot illustrates the expression levels of these markers, where the color intensity of the bubbles indicates expression levels, and the size reflects the relative abundance of each cell type.

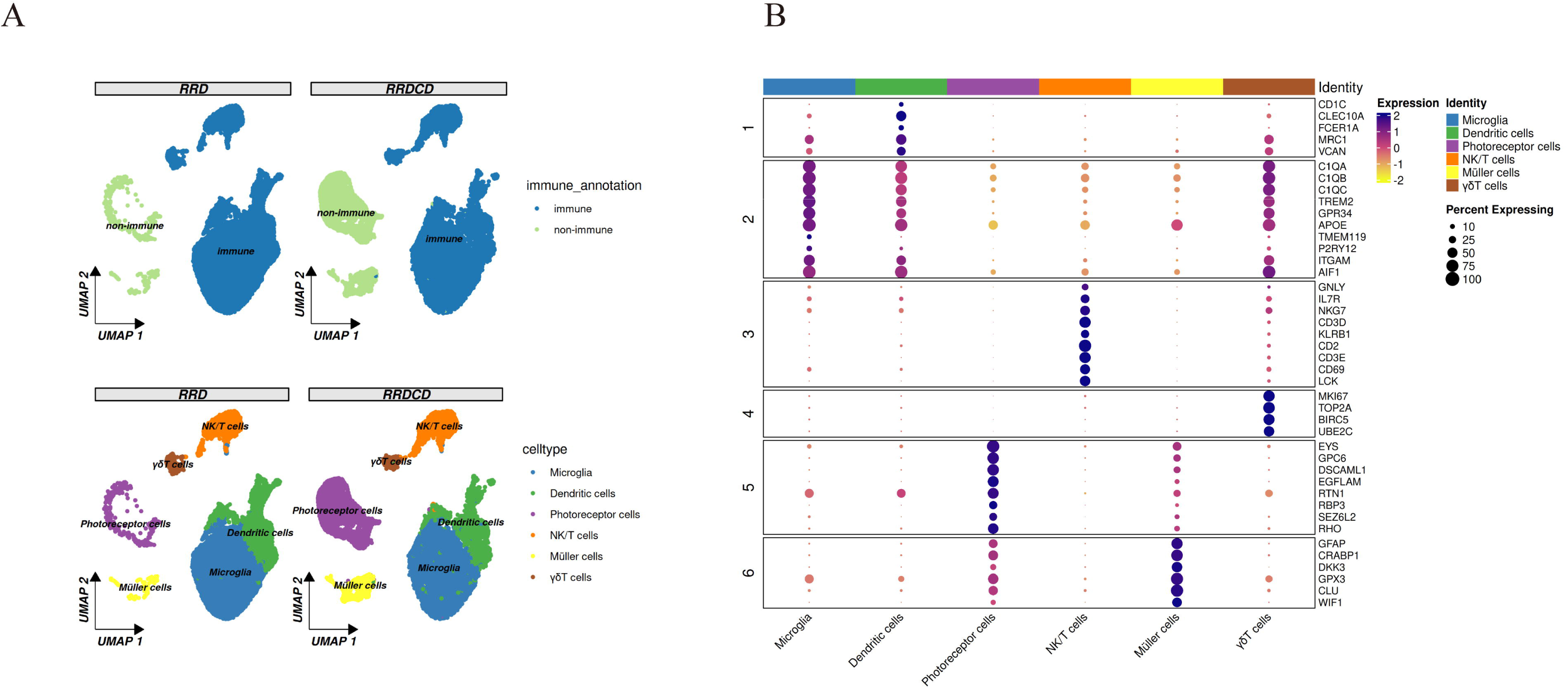

### Intercellular Communication Networks in RRDCD

To investigate differences in retinal cellular interactions between RRD and RRDCD groups, we employed CellPhoneDB to analyze their molecular communication networks. The heatmaps in Figure 3A illustrate these dynamics, with the RRDCD group exhibiting notably stronger connectivity between microglia and dendritic cells. These findings highlight significant differences in molecular interactions between the two groups, emphasizing enhanced crosstalk between microglia and dendritic cells in the RRDCD context.

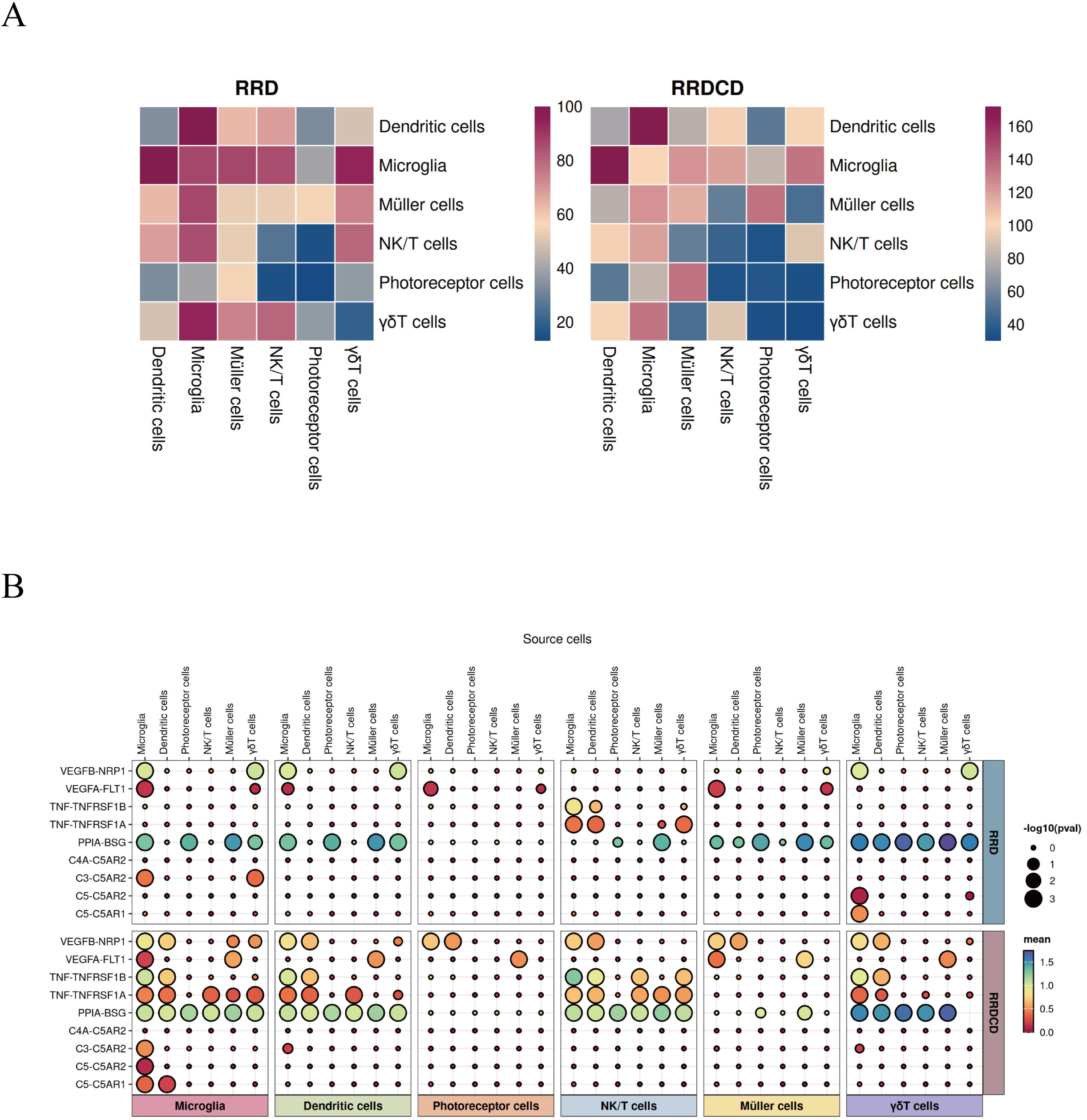

Specifically, the complement C5-C5AR1 interaction was notably enhanced in microglia and dendritic cells of the RRDCD group (Figure 3B), suggesting that these immune cells play a crucial role in modulating inflammatory responses during disease progression. Moreover, interactions involving VEGFA with NRP1 and FTL1 in the RRDCD group showed variability, with some interactions being enhanced while others were reduced between different cell types. Additionally, we identified other significant receptor–ligand interactions, such as complement C4A with C5AR2 and complement C3 with C5AR2, indicating a complex signaling network that modulates immune responses in retinal degeneration. Overall, the enhanced complement C5-C5AR1 signaling may facilitate the recruitment and activation of other immune cells, further amplifying the immune response during retinal degeneration.

### Impact of Complement C5 on Microglial Activity and Endothelial Cells

We assessed the effect of complement C5 on primary mouse retinal microglia and its influence on the activity of RF/6A choroidal vascular endothelial cells. Primary microglia were treated with varying concentrations of complement C5 (0, 0.5, 1.0, 2.5 µg/ml) for 24 hours, followed by co-culture with RF/6A cells. MTT assay results (Figure 4A) demonstrated a dose-dependent reduction in RF/6A cell viability post C5 treatment, indicating that C5-treated microglia negatively affect the survival of endothelial cells. Furthermore, the TUNEL assay (Figure 4A) confirmed that C5-treated microglia induced apoptosis in RF/6A cells, suggesting a potential mechanism through which microglia activation can influence vascular integrity in the retina.

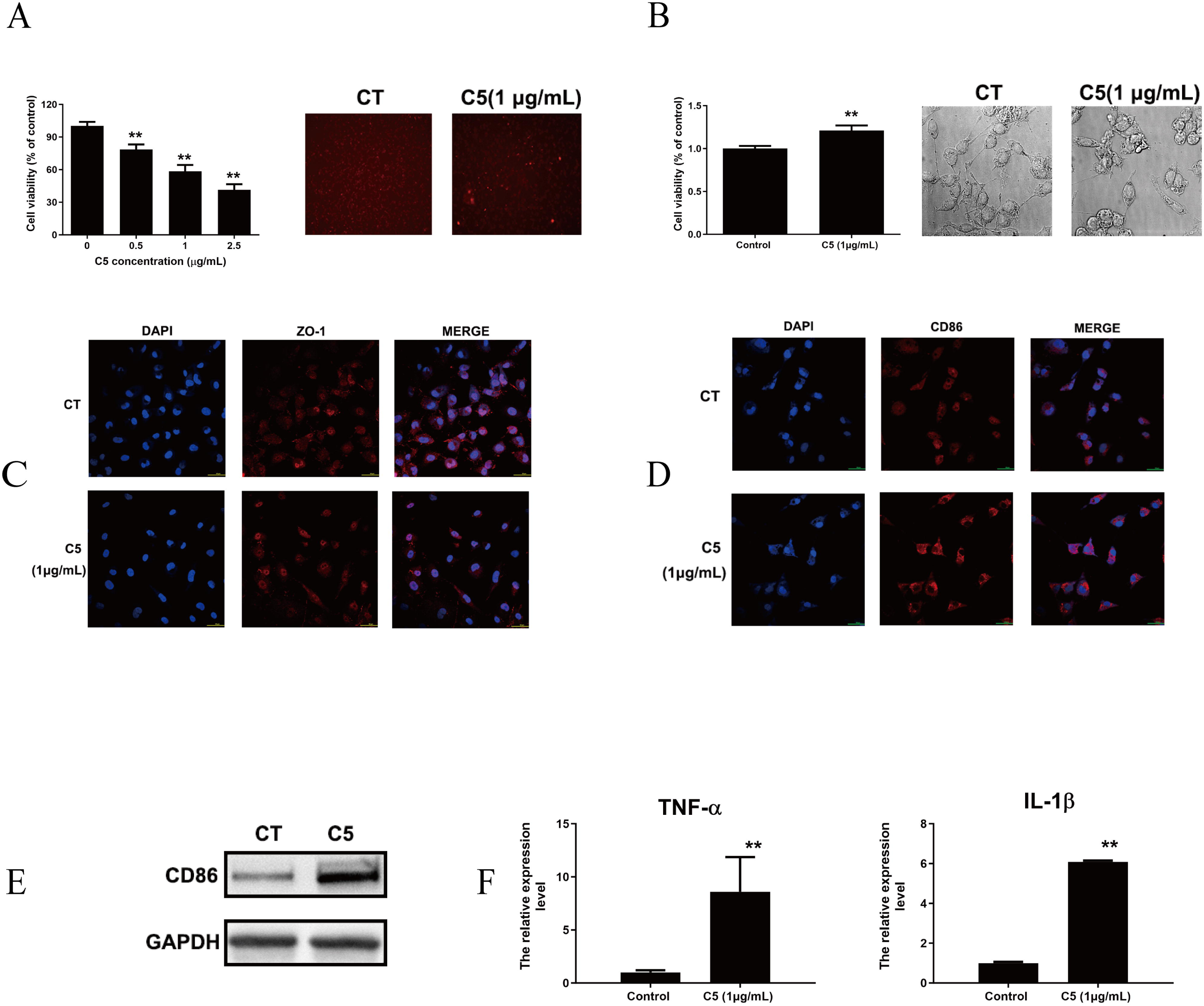

### Complement C5 Promotes Microglial Proliferation

To assess the impact of complement C5 on the proliferation of primary mouse retinal microglia, cells were treated with C5 at a concentration of 1.0 µg/ml for 24 hours. Morphological analysis revealed that C5 treatment significantly stimulated microglial proliferation. Under a phase-contrast microscope, C5-treated cells exhibited increased cell density and enhanced extension of cellular processes, indicative of an activated state. Additionally, the MTT assay (Figure 4B) demonstrated that C5 treatment significantly increased the metabolic activity of microglia compared to the control group (p < 0.05).

### Disruption of Tight Junctions in RPE Cells by C5-Treated Microglia

Given that different concentrations of complement C5 led to RF/6A cell apoptosis, we focused on a concentration of 1.0 µg/ml C5 for subsequent experiments. Microglia were co-cultured with ARPE-19 retinal pigment epithelial cells for 24 hours following treatment with 1.0 µg/ml C5. Immunofluorescence analysis of the tight junction protein ZO-1 revealed significant disruption in the localization and intensity of ZO-1 in ARPE-19 cells exposed to C5-treated microglia (Figure 4C). This finding indicates that complement C5 treatment leads to the impairment of tight junctions in RPE cells, potentially contributing to retinal barrier dysfunction.

### Pro-Inflammatory Responses Induced by Complement C5

To investigate the inflammatory effects of complement C5 on microglia, we examined the expression of CD86, a marker of M1 polarization, via immunofluorescence and Western blot analysis after 24-hour treatment with 1.0 µg/ml C5. Results showed a marked increase in CD86 expression in C5-treated microglia (Figure 4D and 4E). Additionally, ELISA assays measuring cytokine levels revealed that complement C5 significantly elevated the expression of pro-inflammatory cytokines TNF-α and IL-1β in microglia compared to controls (Figure 4F). These findings demonstrate that complement C5 can potentiate the inflammatory response in retinal microglia, leading to increased cytokine release and suggesting a role in retinal inflammation and associated pathology.

## Discussion

In this study, microglia emerged as the predominant cell type in vitreous samples from both RRD and RRDCD patients. Clinically, photoreceptor cells (PCs) associated with RRD were detected only in small quantities in vitreous samples, likely because single-cell RNA sequencing detects only live cells, and free PCs in RRD may have already died. This results in the predominance of active microglia and a reduced number of PCs. Our cell–cell communication analysis revealed that the RRDCD group exhibited notably stronger connectivity between microglia and dendritic cells compared to the RRD group. Among the significant interactions, the complement C5-C5AR1 pathway was prominently enhanced in microglia and dendritic cells of RRDCD patients. This suggests that the complement C5/C5a signaling axis plays a critical role in modulating inflammatory responses during the progression of RRDCD. Additionally, interactions involving VEGFA with NRP1 and FTL1 showed variability, indicating a complex network of signaling pathways that contribute to the disease’s inflammatory milieu.

Our in vitro experiments further elucidated the impact of complement C5/C5a on microglial activity and its downstream effects on endothelial and epithelial cells. Treatment of primary mouse retinal microglia with complement C5 led to increased microglial proliferation and activation, as evidenced by elevated metabolic activity and enhanced extension of cellular processes. These activated microglia, when co-cultured with RF/6A endothelial cells, induced a dose-dependent reduction in cell viability and increased apoptosis, highlighting the detrimental effects of microglial activation on vascular integrity in the retina. Moreover, co-culture of C5-treated microglia with ARPE-19 retinal pigment epithelial cells resulted in significant disruption of the tight junction protein ZO-1. This impairment of tight junctions suggests a compromise in the blood–retina barrier, which is critical for maintaining retinal homeostasis. The degradation of tight junctions can facilitate the leakage of choroidal blood vessels, contributing to the choroidal detachment observed in RRDCD.

Our findings also demonstrate that complement C5 significantly elevates the expression of pro-inflammatory cytokines TNF-α and IL-1β in microglia. The increased cytokine release underscores the role of complement C5/C5a in potentiating the inflammatory response within the retinal environment. This heightened inflammation not only exacerbates retinal damage but also creates a feedback loop that further activates the complement system, perpetuating the cycle of inflammation and tissue disruption.

Previous studies have established that complement-related signaling axes, such as C3a-C3aR and C5a-C5aR, are pivotal in triggering and regulating inflammatory responses, and are associated with numerous inflammation-related diseases ^[14,15]^. Consistent with these findings, we identified complement C5a-related receptor-ligand interactions in the vitreous of both RRD and RRDCD patients, with a more pronounced presence in RRDCD. This aligns with the clinical observation that RRDCD is characterized by more severe inflammatory responses compared to RRD.

Regarding the pathogenesis of RRDCD, two main hypotheses have been proposed: one emphasizes a hypotonous origin ^[16]^, while the other stresses an inflammatory origin ^[2]^. Our single-cell sequencing analysis revealed that both RRD and RRDCD vitreous fluids are predominantly composed of microglia. Furthermore, cell–cell communication analysis indicated that ligand-receptor interactions in both groups are primarily related to the blood–retina barrier and complement C5a, with most interactions occurring between microglia and other cell types. This suggests that an inflammatory response is already present in RRD, which is further intensified in RRDCD. Our in vitro experiments confirmed that complement C5/C5a stimulation of microglia can disrupt the blood–retina barrier and induce choroidal leakage, supporting the notion that complement C5a in RRD contributes to the exacerbation of blood–retina barrier disruption and choroidal leakage in RRDCD. Consequently, our findings favor the hypothesis that inflammation is the primary driver (“cause”) of RRDCD development.

Furthermore, the regulation of retinal microglia involves intricate interactions with the retinal microenvironment. Recent cell fate mapping studies confirm that adult retinal microglia exist as a self-maintaining closed population under healthy conditions, with their numbers, distribution, and physiological states being highly organized and stable ^[17–19]^. Microglial homeostasis is tightly regulated by signals from surrounding neurons and other retinal cells, including cytokines, chemokines, complement components, and damage-associated molecular patterns ^[20]^. For instance, TGFβ secreted by retinal pigment epithelial (RPE) cells can induce microglia to release IL-10, downregulating the expression of antigen-presenting molecules such as MHC-II, CD80, and CD86, thereby promoting an anti-inflammatory phenotype ^[21]^. Additionally, direct physical interactions between microglia and other retinal cells, mediated by molecules like CD200-CD200R and CX3CL1-CX3CR1, play crucial roles in inhibiting pro-inflammatory activation and maintaining microglial surveillance capabilities^[22–26]^.

The relationship between microglia and the complement system is intricate. Microglia express complement receptors, ligands, and activation factors, and are involved in processes such as synaptic pruning through the complement receptor 3 (CR3) complex ^[27,28]^. In addition to being effectors of complement-mediated phagocytosis, microglia themselves are significant sources of complement components in the retina. Our single-cell sequencing data corroborate these findings, showing a close association between microglia and complement C5 in RRDCD. In vitro experiments demonstrated that complement C5/C5a interaction with microglia enhances their pro-inflammatory state, leading to blood–retina barrier disruption and choroidal leakage.

Overall, the current study supports the inflammatory hypothesis of RRDCD pathogenesis, highlighting the pivotal roles of microglia and the complement C5/C5a pathway in exacerbating retinal inflammation and compromising vascular integrity. However, the study has limitations, including the lack of human retinal microglia for in vitro studies and a limited sample size due to the COVID-19 pandemic in China. Future research should aim to include larger patient cohorts and employ rigorous animal models to further elucidate the mechanisms by which microglia and complement components contribute to RRDCD. Such studies are essential for developing targeted therapeutic strategies aimed at mitigating inflammation, preserving the blood–retina barrier, and improving surgical outcomes for patients suffering from this debilitating condition.

## Acknowledgments

Huiyan Xu and Qiuhong Wang contribute equally to this work and are co-first authors.

